# Modelling conduction delays in the corpus callosum using MRI-measured g-ratio

**DOI:** 10.1101/479881

**Authors:** S. Berman, S. Filo, A. A. Mezer

## Abstract

Conduction of action potentials along myelinated axons is affected by their structural features, such as the axonal g-ratio, the ratio between the inner and outer diameters of the myelin sheath surrounding the axon. The effect of g-ratio variance on conduction properties has been quantitatively evaluated using single-axon models. It has recently become possible to estimate a g-ratio weighted measurement *in vivo* using quantitative MRI. Nevertheless, it is still unclear whether the variance in the g-ratio in the healthy human brain leads to significant differences in conduction velocity. In this work we tested whether the g-ratio MRI measurement can be used to predict conduction delays in the corpus callosum.

We present a novel framework in which the structural properties of fibers (i.e. length and g-ratio, measured using MRI), are incorporated in a biophysical model of axon conduction, to predict conduction delays of long-range white matter fibers. We applied this framework to the corpus callosum, and found conduction delay estimates that are compatible with previously estimated values of conduction delays. We account for the variance in the velocity given the axon diameter distribution in the splenium, mid-body and genu, to further compare the fibers within the corpus callosum.

Conduction delays have been suggested to increase with age. Therefore, we investigated whether there are differences in the g-ratio and the fiber length between young and old adults, and whether this leads to a difference in conduction speed and delays. We found small but significant differences between the predicted delays of the two groups in the motor fibers of the corpus callosum. We also found that the motor fibers of the corpus callosum have the fastest conduction estimates. Using the axon diameter distributions, we found that the occipital fibers have the slowest estimations, while the frontal and motor fiber tracts have similar estimates.

Our study provides a framework for predicting conduction latencies *in vivo*. The framework could have major implications for future studies of white matter diseases and large range network computations. Our results highlight the need for improving additional *in vivo* measurements of white matter microstructure.

## 1. Introduction

White matter tissue consists of myelinated and non-myelinated axons and glial cells, forming large-scale networks which are necessary for learning (1–3), cognitive functions (4), normal development, and normal aging (5,6). Furthermore, damage to the structure of the white matter has been associated with a variety of diseases and disorders, including multiple sclerosis and schizophrenia (7,8). These networks of white matter fibers are essential for brain function because their main role is to conduct action potentials between distant brain regions. The conduction delays, defines as the time required for information to travel along white matter fibers, could have computational implications. The conduction delays were also shown to vary with many factors, including age (9), sex (10), and white matter diseases such as multiple sclerosis (11).

*In vivo* anatomical studies of white matter often use indirect measures of its structure and integrity properties, which would benefit from a biophysical framework to relate them to white matter function (12–14). *Ex vivo* and *in vitro* studies have led to a rich understanding of the biophysical connection between the axon microstructure and its conduction properties (e.g., (15–17)). Moreover, computational models of conduction of action potentials along myelinated axons provide a way to explore the dependence of the conduction properties on the axons’ geometrical and electrical properties.

One such geometrical property of axons is the g-ratio (*g*). It is defined as the ratio between the inner and outer diameters of the myelin sheath surrounding the axon. For a fixed inner diameter (axon diameter, *d*), changing *g* means changing the myelin sheath thickness, thus affecting membrane resistance and capacitance. For a fixed outer diameter (fiber diameter, *D*), changing *g* means varying both myelin sheath thickness and intra-axonal space, thus affecting axial conductivity as well. Under these assumptions Rushton found analytically that in the peripheral nervous system, an optimal *g* for conduction velocity is around 0.6 (18). Further work, modelling the central nervous system of the rat, has shown that higher values (~0.77) would be optimal given other constraints on the tissue such as energy consumption or volume limitations (19).

Empirical values of g-ratio can be measured directly using invasive techniques such as electron microscopy. Such studies reveal the *g* is indeed distributed around near optimal values: the reported values are around 0.67 for monkey’s corpus callosum (20,21), 0.81 for guinea pigs’ optic nerve (22), 0.76 for the mouse corpus callosum (23) and 0.75-0.8 for the rat corpus callosum (24). It has recently become possible to estimate a *g*-weighted measurement in humans *in vivo* using quantitative MRI (qMRI) (25–29). These measurements provide values and trends that are similar to those found with histology (21,27). Nevertheless, it is unclear whether the variance captured by the *in vivo g* measurements in the healthy human brain can be translated to meaningful differences in conduction velocity.

Theoretical estimates of conduction time across the corpus callosum have been previously used in cross-species studies (30). *In vivo* estimations of callosal delays can be done using electrical recordings such as EEG or MEG. Such measurements have been shown to be related to callosal structure (31,32), and are likely to increase with age (33,34). The increase in callosal delay time with age could be related to white matter structure, as it has been shown that during aging the brain tissue structure undergoes changes, with accumulating damage to the white matter (35,36), and a decrease in myelination (37). It is still an open question whether *g*-weighted MRI can be related to age dependent changes in callosal delay time.

Both theoretical work, and *in vivo* studies (mentioned above) show a relationship between indirect measurements of white matte structure and function, yet it is important to establish this relationship empirically. Animal studies that provide estimates of both axon geometry and conduction velocity are rare, particularly in mammalian central nervous system. Nevertheless, there is some empirical evidence for the relationship between axon diameter, myelin thickness and conduction velocity. Hursh used excised nerves of the cat to display a linear relationship between axon diameter and conduction velocity (38). A similar dependence was found on the frog sciatic nerve (39). Furthermore, a strong relationship was found between myelin thickness and conduction velocity in the rabbit peroneal nerve (40).

The results of the animal studies are in agreement with the theoretical work, and both have existed for dozens of years, highlighting a great need for understanding the functional meaning of structural properties of white matter. Recent advances in qMRI measurements allow to estimate white matter microstructure *in vivo*. Importantly, it is still unclear how the structural measurements done with MRI can be biophysically related to conduction along the white matter fibers. To test whether the *g*-weighted MRI measurements can be used to model callosal delays, we first discuss a theoretical framework that can be used to calculate conduction velocity for a single axon. Next, we describe how we implement this model to calculate conduction delays in the corpus callosum *in vivo* with MRI. This is the first time a framework is presented that related MRI measurements in white matter to conduction properties using biophysical models of conduction in myelinated axons. Finally, we use our framework to estimate conduction in both younger (under 40) and older (over 65) subjects. Thus, we test whether incorporating the *g*-weighted MRI measurement in a biophysical model of conduction allows us to explain the previously observed increase in delay as function of age.

## 2. Theory – predicting conduction latencies

In the theory section we propose a framework that relates MRI measurements of averages of microstructure features to single axon conduction models, in order to predict white matter conduction *in vivo*. For this purpose, we first identify a relevant axon model, and the essential parameters in the model (section 2.1). Then, we describe the available qMRI parameters that can be used to estimate those model parameters (section 2.2). For model parameters that cannot be estimated based on qMRI measurements, specific values must be assumed. We will describe the assumptions underlying the choice of specific values for those parameters (section 2.3). In this section we also discuss how these parameters can be applied to a model for predicting conduction along a certain tract.

### 2.1 Conduction in myelinated axons

The myelinated fiber is comprised of two qualitatively different sections: the nodes of Ranvier and the internodes. The nodes of Ranvier, which contain a high density of voltage-gated Na+ channels can generate action potentials. The internodes are long sections between the nodes (about 10^3^ longer), and they are coated by many layers of myelin sheaths (41). Due to the myelin sheath, the voltage progression in the internode can be described as passive. The progression of voltage along both segments of the axons have already been modelled and numerically simulated (e.g., 12,24–26). This understanding of the axon signal conduction, together with knowledge on the axon’s electrical properties (45,46) and the ability to measure its geometry, will allow to implement a model that links white matter structure with function in biophysical terms.

#### 2.1.1 Dependence of conduction velocity on axon geometry

The determinants of conduction velocity of action potential propagation along a myelinated axon have been previously described and modelled. Rushton (18) was the first to describe the dependence of the conduction velocity of a myelinated axon on its geometry. Rushton showed that for a myelinated axon, the conduction velocity (*θ*) is proportional to internode length (*l*). The internode length, in turn, is roughly proportional to the fiber diameter (*D*), which can be defined using the axon diameter (*d*) and the g-ratio (*g*): *D* = *d*/*g*. According to his formulation, we could predict conduction velocity, up to a proportionality factor, using the axon diameter and *g* (equation 1): 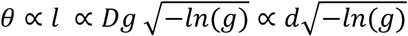 Rushton’s formulation provides an expression proportional to conduction velocity. Following studies have replicated Rushton’s predictions and shown the dependence of the axon’s conduction velocity on its diameter, internode length, and myelin sheath thickness (47–50). Others studies evaluated the effect of additional parameters of the axon on conduction velocity. For example, Moore et al. (42), using numerical simulations of myelinated fibers, examined both geometrical and physiological parameters. The simulations replicated the results showing a dependence of conduction velocity on internode properties, while also revealing that node parameters have a minimal effect on the conduction velocity.

Moore’s approach of numerically simulating the voltage propagation, provides an approximation of conduction velocity, and allows a ‘direct’ manipulation of the different axon properties, which could be useful for the purpose of this study. In this study we use a recent MATLAB implementation that numerically simulates the conduction along a myelinated axon using Richardson’s axon model-C, and Halter and Clark’s numerical formalization (43,51,52). Specifically, the simulation models the axon by dividing it into compartments representing the node, paranode and internode. For each time step, current flows are calculated. The current flows across the membrane (either the axonal membrane or the total myelin membrane) are calculated from the values of voltage, voltage change rate, and the specific membrane capacitance and conductance. The intracellular axial current flow is calculated from the specific intracellular resistance and the gradient of intracellular voltage. In the simulations, each model axon was subjected to a current stimulation of amplitude 3nA applied for 10ms to the first node. The conduction velocity was then measured over a 10-node interval between the 20th and 30th node. The simulations were checked to ensure membrane potential peak of at least -20 mV were achieved on a minimum of 10 consecutive nodes.

### 2.2 The measurements in qMRI

Using a biophysical model, such as the simulation described here, allows for the computation of the conduction time of an action potential along a “theoretical axon”, which represents an average axon in an axonal fascicle. In order to do so *in vivo*, we should be able to measure white matter pathways and estimate their microstructure. In particular, in the section below we describe ways to measure the necessary geometrical measurements: the axon diameter, *g*-ratio, and axon length. In the following subsections, we describe recent advances in qMRI, which allow for ongoing improvement in estimating the axon diameter, the g-ratio and the overall tract length.

#### 2.2.1 Diffusion and axon properties

Diffusion MRI (dMRI) provides an important tool for measuring white matter microstructure *in vivo*. Water molecules impinge on cellular membranes, intracellular organelles, neurofilaments, and myelin. The diffusion displacement distribution therefore contains information about all of these structures. When the diffusion is measured in different directions, models can be applied to the data, and these models’ parameters can be used to represent the white matter microstructure (53).

##### Axon diameter

There are several methods of acquiring and modelling dMRI to measure the mean axon diameter or the axon diameter distribution. Currently most of the methods are challenging since they require multi-shell high angular resolution diffusion imaging (HARDI) data, and special diffusion acquisition sequences that use either very strong and/or oscillation gradients. Furthermore, the dMRI resolution limit is very far from the resolution of axon bundles, and current methods have a limited sensitivity for small axons (54). The models used for estimating axon diameter *in vivo* often have many assumptions on tissue properties, that might be invalid, or require knowledge of the main orientation of fibers. Due to these limitations, *in vivo* measurements of axon diameter are challenging and some controversy remains regarding their feasibility (55). Nevertheless, some of the existing methods have been validated using histology and phantoms of cylinders in varying diameters (54,56–59).

##### g-ratio

The voxel average g-ratio may be estimated using a model combining dMRI parameters reflecting axon or fiber volume in the voxel (Fiber Volume Fraction, FVF), with an MRI-myelin measurement (Myelin Volume fraction, MVF) (25). The dMRI models that were previously used to calculate g-ratio include neurite orientation dispersion and density imaging (NODDI) (21,60), ActiveAx (28,61), AxCaliber (28,56), tensor fiber density (TFD) (29,62), and tensor modeling (25,63,64). It is important to note that each method has its own advantages and limitations. In many cases the limitations of these estimates come from the assumptions of their models, that might not hold true in tissue with crossing fibers and in tissue with compartments with different diffusion properties. The MVF also saw several implementations including mcDESPOT (26,65), multi-exponential T2 (66–68), T2* (69,70), quantitative magnetization transfer (qMT) (25,29,63,68,71–74), and non-water fraction (27,28,75). Most of the MVF estimates, while sensitive to myelin, are not specific. Therefore, some of these methods require calibration, and/or may provide a poor estimate of MVF in tissue that is not healthy white matter tissue (63,70,76). The myelin water fraction is arguably the only specific MVF MRI measurement (67), yet it is currently hard to measure in clinical scanners which often comes with a reduced signal to noise ratio (SNR). Recent work compared the reliability of several of the different g-ratio methods and found NODDI to be more reliable than FA for the FVF estimation, where MTV and qMT were similar, with qMT having some advantage (77). Despite the limitations above, the MRI measurement of g-ratio was shown to provide a good estimate of the weighted average of the g-ratio in a voxel (78).

##### Length

Tractography algorithms use dMRI to reconstruct the white matter fiber tracts (79,80). The tract reconstruction provides the pathway length which is a critical parameter for deriving conduction time from conduction velocity estimates.

##### Tract properties

Since this work aims at assessing conduction in white matter, it is necessary to characterize the white matter microstructure properties of specific relevant white matter fibers. Using tractography one can measure structural MRI measurements (*g*, *d*), along the reconstructed white matter tracts, either along the core (i.e. the mean position of the streamlines’ position per node), per voxel, or per streamline (79,81,82).

#### 2.2.2 Modelling Local Field Potential (LFP) using MRI data

Having performed tractography, and measuring g-ratio and axon diameter along relevant fiber tracts, one can then use these measurements in the axon model and predict the mean conduction time for an individual or groups of subjects. Additional analysis of the data might reveal more about conduction properties. For example, one can use the variance of estimated latencies within a single subject’s white matter tract, and try to model the local field potential. The framework implemented in a recent study (83) allowed for a simple model that simulates LFP measures. Modelling an LFP from the MRI data allows for a more comprehensive comparison with the electrical signal: ERP width, amplitude etc. While these properties are also likely to be affected by cortical processing as well as skull properties, it is worth testing the extent to which the white matter contributes to the recorded electrical signal.

### 2.3 Simulation and choice of parameters

Several of the model parameters cannot be measured with MRI, and must be set manually. These parameters include the axon geometry, and specific electrical properties of the axon (e.g. specific axial resistance). Importantly, the axon’s specific electrical properties have been measured in rodents and primates, and are considered to be conserved over species (46). Nevertheless, exceptions are possible, as evident from a recent study which found that the human membrane capacitance is 0.45 µ*F*/*cm*^2^ and not 0.9 as is the case with mice (84).

While the electrical properties of the axon are thought to be conserved, the geometrical aspects have been shown to vary with age, disease, and possibly with learning and behavior. Therefore, a potential limitation in modeling the human white matter conduction time is that not all geometrical parameters can be measured *in vivo*. These parameters will have to be based on the literature. The internode length, for example, is a significant parameter (48,85–87), yet MRI measurements do not allow such information to be extracted. Fortunately, many geometrical properties of the axon share a close to linear relationship: It has been shown that *g* can be described as a monotonic function of the axon diameter, and that the internode length is roughly a linear function of the axon diameter (16,24,38,88). The latter relationship was used to reach Rushton’s formulation, and it can be used explicitly when assuming an internode length for the simulation.

We explored the conduction velocity as calculated with the simulation, for a range of values for axon diameter (*d*) and fiber diameter (*D*)). We chose values that were found for myelinated axon in the human central nervous system (89), with values ranging from 0.2 to 5 *µm* for both *d* and *D* The internode length was calculated as a linear function of the fiber diameter, as found in the anterior medullary velum of the adult rat: *L* = 117 + 30 · *D* (90). The specific membrane capacitance was set to 0.45 µ*F*/*cm*^2^ (84). The rest of the (electrical) properties were fixed as described in table 1 of Lorena et al,’s paper (52). It can be seen in Figure 1a that the larger the fiber diameter (*D*), the faster it will conduct. Furthermore, for a given fiber diameter, the conduction velocity as a function of axon diameter (*d*) has a maximum around *g*=0.6, as shown by Rushton. The intuition for this is the following: given a constant fiber diameter, dramatically decreasing *g* means a very small axon diameter, thus reducing intra-axonal conductivity resulting in slower conduction. However, dramatically increasing *g* will reduce the leak resistance (46), which will result in slower conduction as well. Therefore, the optimal *g* should lie somewhere in the middle, and Rushton’s analysis found it to be 0.6. As mentioned in the Introduction, Rushton’s analysis considered the peripheral nervous system. More recent theoretical study focused on the rat central nervous system (19), and found that given volume and energy consumption constraints, the optimal *g* should be approximately 0.77.

**Figure 1:**
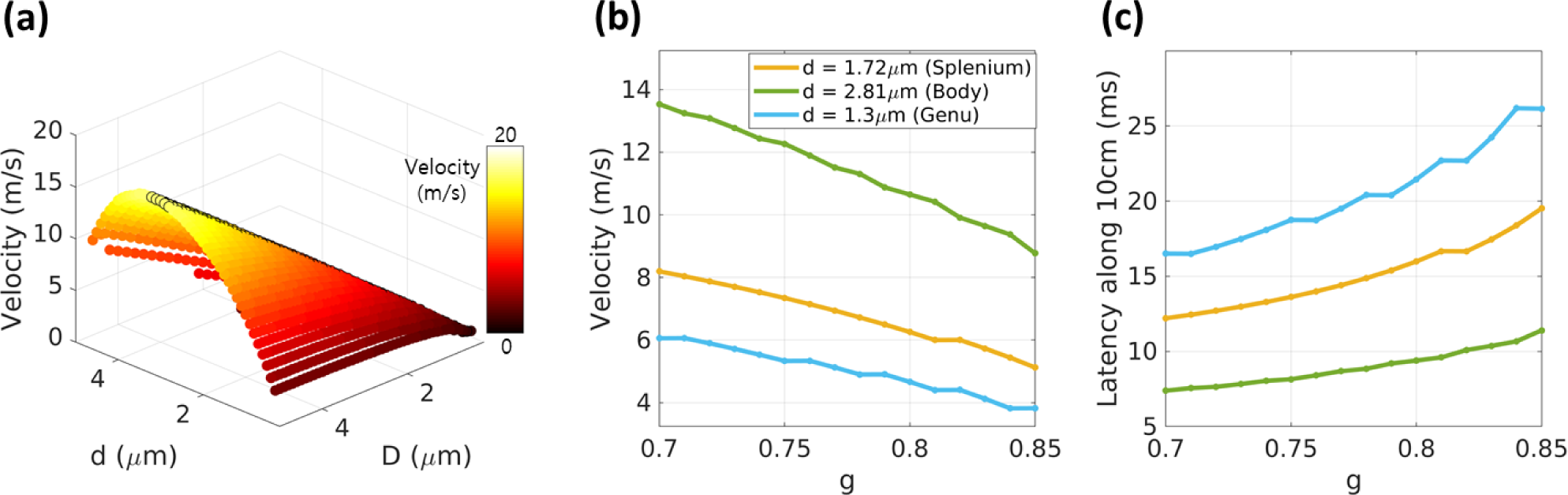
Simulating conduction as a function of axonal properties. A numerical simulation (51) for estimating the conduction velocity. **(a)** Conduction velocity plotted as a function of axon (d) and fiber (axon + myelin) diameter (D), peak velocity can be found around g-ratio=0.6, marked with black circles. **(b)** Conduction velocity as a function of a small biological range of *g*, plotted for three values of axon diameter (1.72,2.81,1.3 *µm*), corresponding to the volume-weighted averages of the axon diameter distributions measured by Aboitiz et al., (1992) in the callosal splenium, mid-body and genu (respectively). **(c)** Conduction delay for the velocities in (b) and a distance of 10cm. The delays for the larger axons are around 5-25ms, similar to inter hemispheric transfer time measured with EEG. The difference in axon diameter between the tracts leads to a large difference in latency.

In the simulation described above, we varied the g-ratio and the axon diameter. However, since in our MRI dataset (as in many others) we cannot calculate an estimate of axon diameter, we test whether measuring *g* alone will be meaningful. To calculate conduction velocity with no estimate of axon diameter, we must first choose and fix the axon diameter for all tracts and subjects. The choice of axon diameter should be informed by known histological measurements, and the fact that MRI measurements, and the g-ratio in particular, are volume-sensitive (91). Aboitiz (89) measured the axon diameter distributions in the human corpus callosum. The results show that the axon diameter distributions are mostly centered around values close to 1*µm*, with very few axons as large as 5*µm* in diameter. To approach the volume-sensitive g-ratio MRI measurement, we calculated a weighted average of the distributions derived by Aboitiz (weighted by axon diameter to give more weight to larger diameters). We find that the weighted average of the callosal splenium, mid-body and genu are 1.72, 2.81 and 1.3 *µm*, respectively. This anatomical segmentation of the corpus callosum corresponds to the functional segmentation of the corpus callosum to its occipital, motor and frontal tracts, respectively.

We predicted velocities of axons with a radius of 1.72, 2.81 and 1.3 *µm* (the weighted average of the callosal axons), given a biologically relevant range of *g*, between 0.7 and 0.85 (Fig. 1b). Next, we estimate the theoretical latency of these axons if they were 10cm long, about the length of motor callosal fibers (Fig. 1c). We find that using our estimates of volume-weighted average of axon diameter in the numerical simulation leads to callosal delays around 7-25 ms. This range is close to that found in electrophysiological studies (92–96), and using response-time differences (33,97). For the following sections, we will use an axon diameter of 1.72, 2.81 and 1.3*µm*, for the occipital, motor and frontal tracts, respectively. It is noteworthy to mention that a smaller axon diameter creates longer delays, and it also increases the effect of *g* on the conduction time estimate (because the velocity is much smaller). Since the motor tract has the largest weighted average of axon diameter, it will most likely have the shortest conduction delay. The conduction delay time in the motor fiber tract won’t be the shortest only if the fiber is substantially longer, or has a dramatically lower g-ratio value. Neither of which are to be expected. For further testing regarding the axon diameter distribution’s effect on the conduction see discussion and Fgure 6.

## 3. Methods

Having established a framework aimed at relating qMRI data to a biophysical model of conduction, we tested whether it would provide sound estimates of conduction latency along white matter fibers. We calculated the predicted latency of signal transfer along callosal fibers in two subject groups that are expected to have variance between them.

### 3.1 Participants

The MRI measurements were performed on 21 young adults (aged 27 ± 2.1 years, 9 females), and 17 older adults (aged 67.4 ± 6 years, 5 females). The Helsinki Ethics Committee of Hadassah Hospital, Jerusalem, Israel approved the experimental procedure. Written informed consent was obtained from each participant prior to the procedure.

### 3.2 MRI Acquisition

Data was collected on a 3T Siemens MAGNETOM Skyra scanner equipped with a 32-channel head receive-only coil at the ELSC neuroimaging unit at the Hebrew University.

#### Quantitative T1, & MTV mapping

3D Spoiled gradient (Flash) echo images were acquired with different flip angles (α = 4°, 10°, 20° and 30°), TE/TR = 3.34/19 ms. The scan resolution was 1 mm isotropic. For calibration, we acquired an additional spin-echo inversion recovery scan with an echo-planar imaging (EPI) read-out (SEIR-epi). This scan was done with a slab-inversion pulse and spatial-spectral fat suppression. For SEIR-epi, the TE/TR was 49/2920 ms. TI were 200, 400, 1,200, and 2,400 ms. We used 2-mm in-plane resolution with a slice thickness of 3 mm. Both the flash and the SEIR-epi scans were performed using 2× GRAPPA acceleration.

#### Diffusion mapping

Whole-brain DWI measurements were performed using a diffusion-weighted spin-echo EPI sequence, accelerated by a factor of 2 (GRAPPA), with isotropic 1.5-mm resolution. The acquisition included 2 diffusion-weightings, one with 32 non-collinear directions (*b*-value = 1000 s/mm^2^) and a second with 64 non-collinear directions (*b*-value = 2000 s/mm^2^). The scan also contained 8 volumes without diffusion weighting (*b*-value = 0). In addition, we collected one scan with six non-diffusion-weighted volumes and a reversed phase encoding direction (posterior-anterior) to correct for echo-planar imaging distortions due to inhomogeneities in the magnetic field.

### 3.3 Estimation of qMRI parameters

#### Quantitative MTV & T1 mapping

Whole-brain MTV and T1 maps, together with bias correction maps of B1+ and B1-, were computed as described in (75,98). In short, unbiased T1 maps were calculated using the variable flip angles which were corrected for B1 excite inhomogeneity using the unbiased SEIR data (99). Next, the T1 maps were used to calculate unbiased proton density (PD) maps. To separate PD from receive-coil inhomogeneity, we assume smooth coil functions and use a biophysical regularization, which finds local linear relationships between 1/T1 and PD. This method was found to be effective and robust to noise (100). The PD was normalized according to values in CSF-only voxels in the ventricles, to produce water-fraction (WF) maps. The MTV maps were then calculated as 1-WF. The analysis pipeline for producing unbiased T1 and MTV maps is an open-source MATLAB code (available at https://github.com/mezera/mrQ).

#### Diffusion parameter estimation

Diffusion analysis was done using the FSL toolbox (101); Susceptibility and eddy current induced distortions were corrected using the reverse phase-encode data, with the eddy and topup commands (102,103). The corrected, unwarped data were aligned to the imaging space of the MTV map using FSL’s Flirt rigid-body alignment (104,105). Furthermore, the data was fit with the Neurite Orientation and Dispersion Diffusion Imaging (NODDI) model using the AMICO toolbox (60,106).

#### g-ratio

In the corpus callosum, the g-ratio was calculated with the following formula, using 2 different measurements: 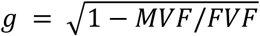. MVF was estimated using MTV (27,28). The FVF was derived from MTV and the isotropic (*V*_*iso*_) and intracellular (*V*_*ic*_) volume fractions estimated from NODDI (21): *FVF* = *MTV* + (1 − *MTV*)(1 − *V*_*iso*_)*V*_*ic*_. Supp. Fig. 1 shows examples of the qMRI maps used in this study (*V*_*ic*_, *V*_*iso*_, *MTV* and g-ratio) for 3 younger and 3 older subjects.

#### Tractography

We performed whole-brain anatomically constrained tractography using the mrTrix software (80). The white matter tracts in corpus callosum were then segmented using the Automated Fiber Quantification (AFQ) toolbox (79). We used this software to track three major callosal tracts connecting the two hemispheres: occipital, motor, and anterior-frontal. We used the AFQ software to sample *g* along the tract core, and create a tract profile of *g* along each tract. The tract profiles are calculated as a weighted sum of each streamline’s *g* value at a given node, where each streamline is weighted based on its Mahalanobis distance from the core of the tract. The result is a vector of 100 equidistant measurements of g-ratio along the trajectory of each callosal tract. The profile can be average over all or some of the nodes to produce a single value for each tract. Furthermore, we use code from the vistasoft git repository to estimate a median *g* value for each streamline. The median value of each streamline is calculated over the 20 nodes around the midsagittal plane. Fig.2 presents an example of the two types of sampling strategies, and Supp. Fig. 2 shows the trajectory of g-ratio along the fiber tracts, averaged across subjects.

**Figure 2:**
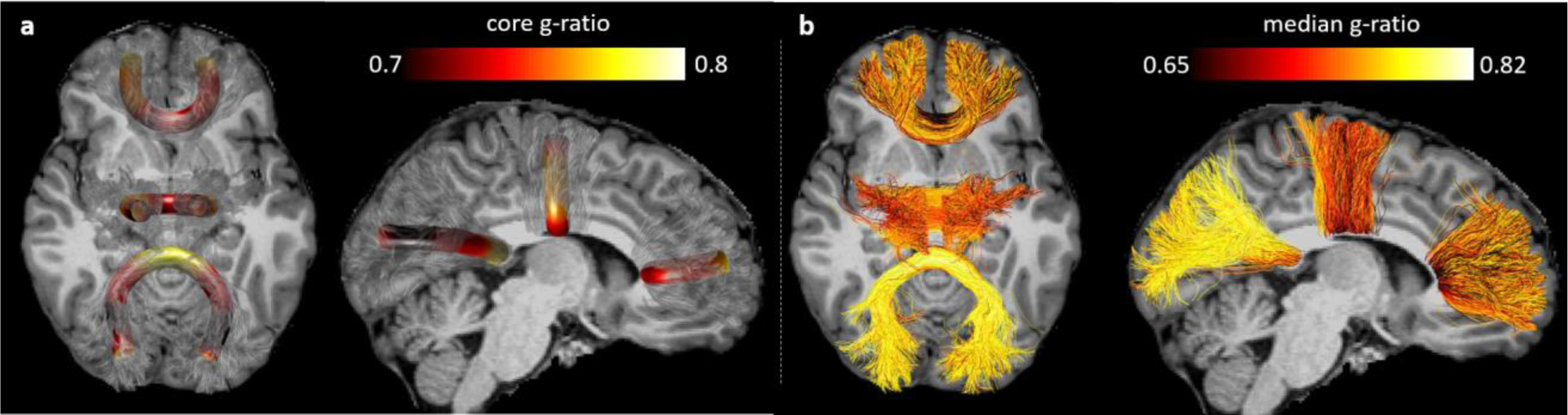
**sampling g-ratio along a fiber tract.** The g-ratio values along 3 callosal tracts of a single subject: the occipital, motor, and anterior-frontal callosal tracts. The g-ratio can either be sampled **(a)** along the tract core, or **(b)** for each streamline separately. The two methods sample the space differently, but both show a variance between and within fiber tracts. The average values of g-ratio along the tract core across subjects can be seen in Supp. Fig. 2.

One young subject was removed from the analysis due to unsuccessful unwarping of the diffusion data. One older subject was removed from the analysis due to extremely sparse tractography results in the occipital cortex. The rest of the subjects (20 young and 16 older subjects) were included in all analyses.

### 3.4 Latency estimation

To test whether we predict a delay in callosal conduction time with age we first calculate the mean of the *g*-weighted MRI measurement along the three callosal tracts. The mean was calculated over the 20 medial nodes of the tract/streamline (for more details on the effect of the choice of node number see Supp. Fig. 2). We incorporate the mean *g* values in the simulation to estimate conduction velocity along the different fiber groups. The model parameters were identical in all simulations with the following exceptions: the g-ratio values were taken from the MRI measurements, the axon diameter was different for each callosal tract (but the same across subjects) and the inter-node length was calculated as a function of g-ratio and axon diameter (see Theory section). The latency of each tract was calculated by dividing the tract length by its velocity. We hypothesize demyelination with age, and a consequent increase in g-ratio, decrease in conduction velocity, and increase in conduction time. Therefore, we used a one-tailed t-test to compare younger and older subjects’ values for each fiber tract, in each of these measures (g-ratio, velocity, and conduction delay).

Finally, we used the calculated latency of each streamline to simulate an LFP signal for each fiber tract. The simulation was adapted from the analysis in (83). The LFP is simulated as the sum of the time-integrated input of a population of neurons. Each neuron receives its input from one streamline. An example of this process is presented in Supp. Fig. 3. First, the median g-ratio value of each streamline is incorporated in the simulation to produce conduction velocity and together with its length, we calculate its conduction delay. The delay times of all streamlines within a single tract, are used as input, which is modeled as Gaussian noise, with a delta function of the corresponding latency of the streamline (Supp. Fig. 3a). Simulating a leaky neuron, the input is integrated over time (Supp. Fig. 3b). Finally, the sum over the streamlines’ simulated time series is treated as the LFP signal. To allow for a comparison between tracts that differ in the number of streamlines, the signal is normalized to have a peak amplitude of 1 [a.u] (Supp, Fig. 3c). The simulated LFP signal has a large peak similar to an event related potential (ERP), and we extracted the peak time and the full width half maximum of the ERP. We used a one-tailed t-test to compare the LFP peak time and two-tailed t-test to compare the width of younger and older subjects’ simulated LFP signal for each fiber tract. Software for the analysis in this study can be found at GitHub [link].

**Figure 3:**
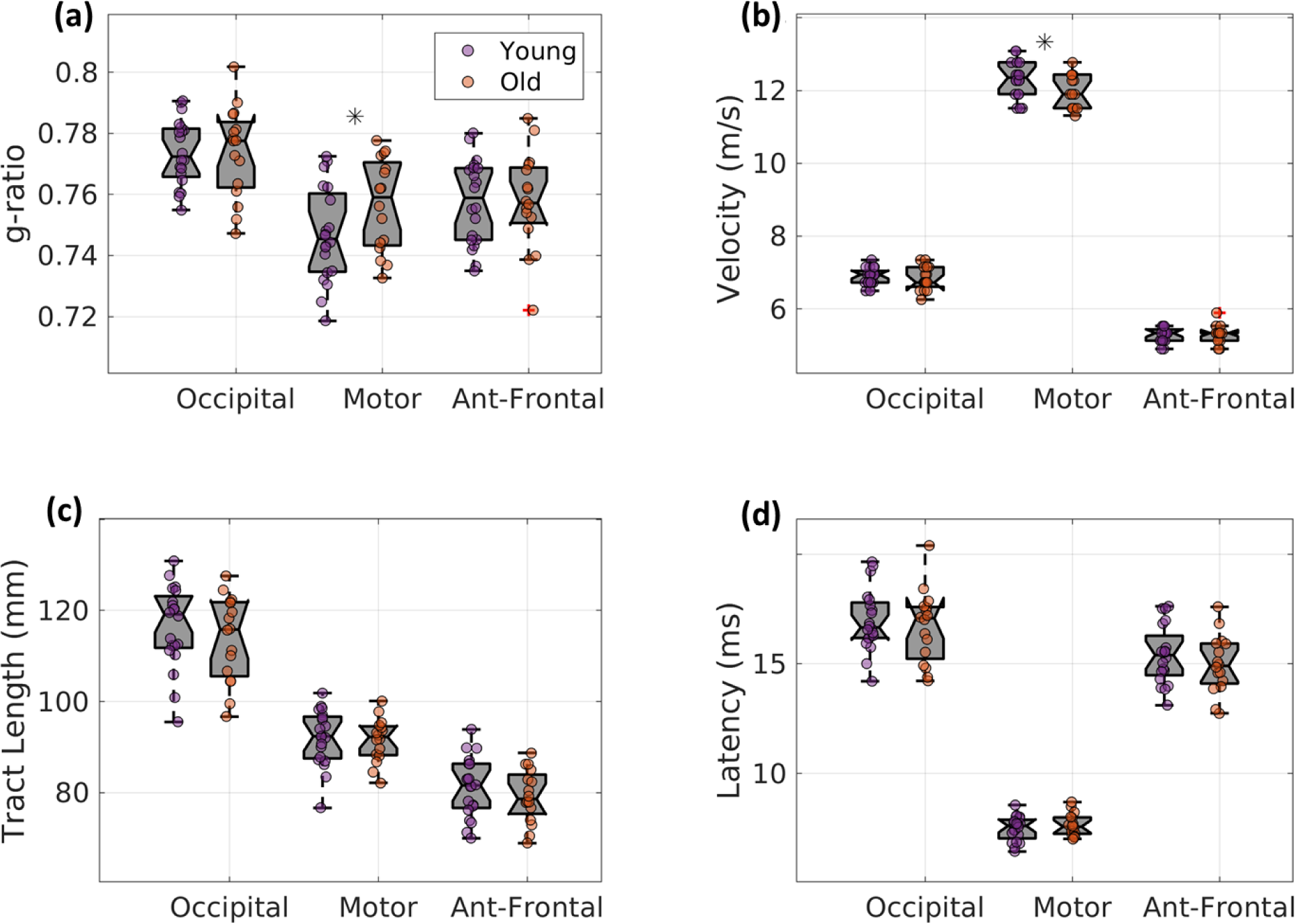
Predicting conduction velocity and latency. **(a)** g-ratio calculated with MTV and NODDI, in three callosal tracts. In the motor callosal region the g-ratio is higher in older (orange) compared with younger (purple) subjects (*p* = 0.03,*t*_32.6_ = −1.96). **(b)** The g-ratio values were used to numerically simulate the conduction velocity along each tract (51). We find that the model predicts a decrease in conduction velocity with age in the motor callosal region (*p* = 0.02,*t*_33_ = 2.13). However, there are no significant differences between the groups’ **(c)** tract length or **(d)** the conduction delays derived from the tract length and the tract conduction velocity.

### 3.5 Axon diameter distributions

Finally, we considered the effect of the known axon diameter distributions on the conduction velocity, and in particular its effect on differences between regions. We reproduced the histograms of the axon diameters in the splenium, genu and mid-body of the corpus callosum, as presented in Figure 4 in Aboitiz et al. (89). To model conduction based on these distributions we performed the same simulations described in the theory and methods section (sections 2.3 and 3.4), with one exception: we estimated the g-ratio for each value of axon diameter using *g* = 0.22 ∙ *log*(*d*) + 0.506 (107). This relationship was derived from axons of the peripheral nervous system, for axons diameter between 0.5*µm* and 4*µm*. Nevertheless, similar trends can be observed in the CNS (78). From the axon diameter and g-ratio distribution we estimated the distribution of velocities of the three regions. We then calculate latency distributions either for a fixed length of 10m or for the mean tract lengths that we measured using the tractography results (section 3.3). Finally, we simulate the LFP of each region, using the distribution of latencies as input to the simulation (sections 2.3 and 3.4).

**Figure 4:**
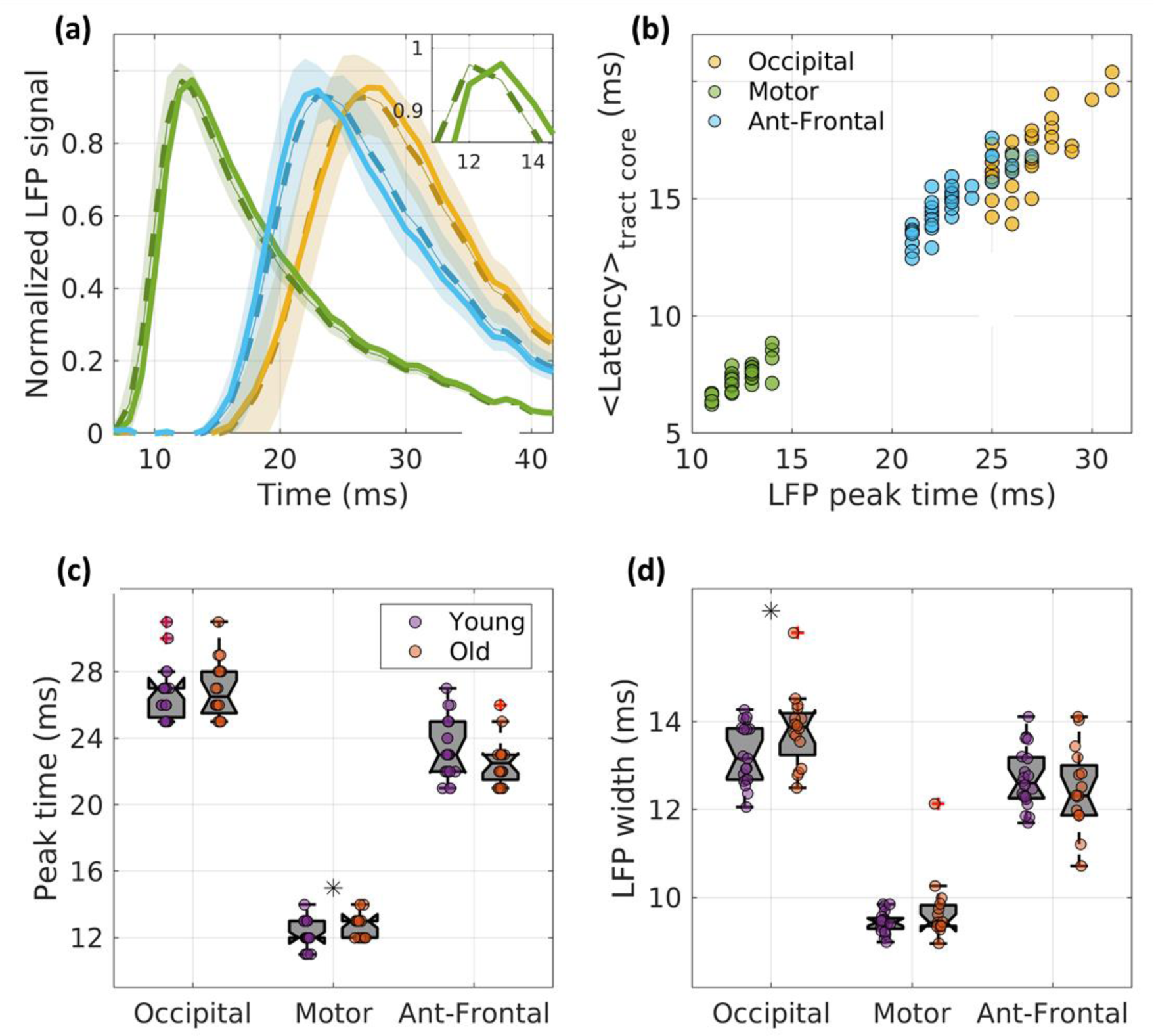
LFP estimates. The g-ratio and length of each streamline was used to calculate the conduction latency. The distribution of latencies for each tract was then used to simulate an LFP signal. **(a)** Simulated LFP for young (solid line) and old (dashed line) in the three fiber tracts. The shaded area is the STD across subjects. The inset zooms in the peak of the LFP for the motor tract of younger and older subjects. **(b)** A comparison of the LFP peak time with the mean latency of the tract core (*R*^2^ = 0.97). The values are not identical but the overall variance is maintained. Group comparison of the LFP peak time **(c)** and width **(d)** reveal different effects than those calculated with the core. The LFP peak time is delayed in the motor callosal region of older subjects (*p* = 0.03,*t*_30.9_ = −1.92). The LFP width of older subjects is significantly larger in the occipital tract (*p* = 0.02,*t*_28.9_ = −2.13).

## 4. Results

To test whether the *g*-weighted MRI measurement can be incorporated in our framework for estimating changes in white matter conduction with age, we first evaluated the *g* values in the corpus callosum of younger and older subjects. Figure 3a shows the average *g* over the 20 medial nodes of the tract core, for each age group and fiber tract (Supp. Fig. 2 shows the effect of the number of nodes taken around the midline). We find that older subjects have higher *g* values compared with younger subjects only in the motor callosal tract (*p* = 0.03, *t*_32.6_ = −1.95, uncorrected for the three comparisons). We also found differences between the callosal regions (see Fig. 5). However, these differences are highly affected by the distance from the mid-sagittal plane suggesting that the calculated g-ratio is sensitive to partial volume effect between white matter tracts (Supp. Fig. 2).

**Figure 5:**
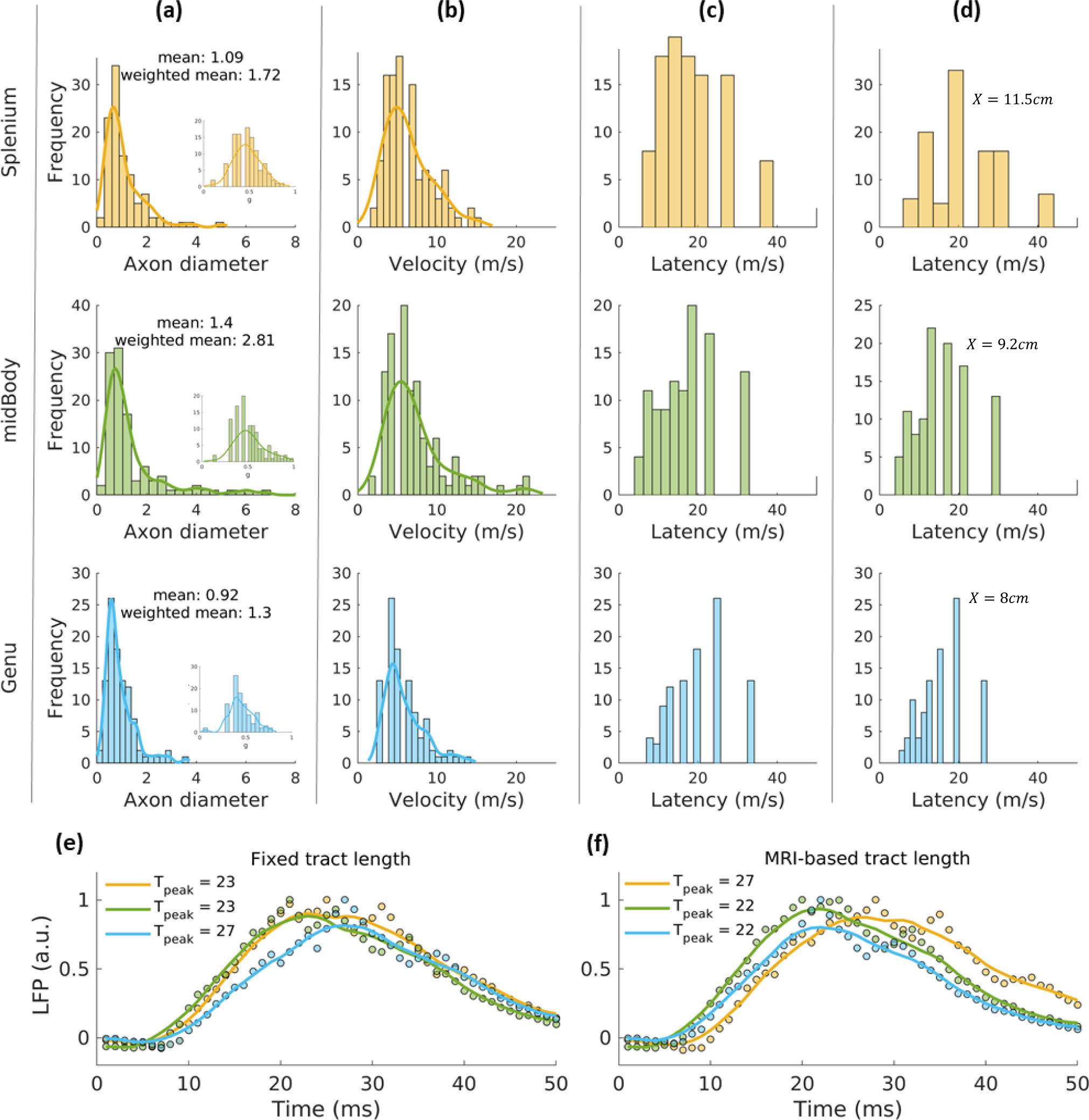
Conduction of axon diameter distribution in the corpus callosum. **(a)** a reproduction of the histograms of the axon diameters in the splenium genu and mid-body of the corpus callosum, as presented in Figure 4 in Aboitiz et al. (1992). **(b)** The distribution of simulated velocities of the three regions, revealing a larger fraction of fast axons in the mid-body. **(c)** The distribution of latencies given a 10cm long axon. **(d)** The distribution of latencies given the tract lengths calculated from tractography: 11.5, 9.2 and 8 cm for the splenium, mid-body, and genu, respectively **(e)** Simulation of the LFP given the latencies in (c), suggests a longer delay in genu fibers. **(f)** Simulation the LFP of each region, given the velocities in (d) and the tract lengths from the tractography data. The results suggest longer conduction delays in the splenium.

We incorporated the *g*-weighted measurements in the simulation to calculate conduction velocities (Fig. 3b). The simulations (Fig. 1b) reveal the dependence of the conduction velocity on g-ratio, (given a fixed axon diameter). The increase of g-ratio with age in the motor callosal region leads to a small but significant decrease of the simulated conduction velocity with age (*p* = 0.02, *t*_33_ = −2.13, uncorrected for the three comparison). Next, we combined the velocity with the fiber tract lengths (Fig. 3c), to calculate conduction time (Fig. 3d). We found no significant differences between the predicted latencies of younger and older subjects, in all areas.

Next, we used the tractography streamlines to predict more conduction properties. We treated each streamline as a separate representative of underlying axonal properties (instead of taking the tract core’s properties to reflect a single value for all streamlines). This sampling approach creates a distribution of *g* values. Each streamline has its own length and g-ratio, thereby resulting in a distribution of estimated latencies. As is for the core (Fig. 3), *g* was sampled along the 20 medial nodes of each streamline, and the median of this sample was used. The distribution of latencies can be used for simulation of an LFP signal (shown in Fig. 4a). The peak time of the LFP is highly correlated with the latency calculated using the tract core (*R*^2^ = 0.97), but it is not identical as it is sensitive to the variance across the tract (Fig. 4b). Comparing the latencies between the age groups (Fig. 4c), we found a delay of conduction latency with age in the motor callosal region (*p* = 0.03, *t*_30.9_ = −1.92, uncorrected for the three comparison).

The LFP analysis also allows us to compare the variance in the data using the signal width. Figure 4d shows the comparison of the full-width half-maximum of the simulated LFP signal between the age groups. Interestingly, this comparison of ERP width between younger and older subjects also reveals a significant difference with age in the callosal fibers connecting the occipital cortices (*p* = 0.02, *t*_28.9_ = −2.13, uncorrected for the three comparisons), suggesting a higher variance within subject, with age. All comparisons between younger and older subjects are not significant after an FDR correction.

Finally, both the core analysis and the LFP simulation predict that the fastest fibers will be in the motor tract of the corpus callosum, with the occipital fibers as much as 10ms slower. To test whether this effect depends on the summary statistic we calculated from the axon diameter distribution, we used the entire distribution to simulate the latency of each callosal regions. The axon diameter distribution shown in Fig. 5a. are adapted from the distribution of the genu, mid-body and splenium, plotted in Fig.4 of Aboitiz’s study (89). Figure 5b displays the distributions of conduction velocities that were simulated from the axon diameter distribution, and their corresponding g-ratio distributions (insets in Fig. 5a). The simulation derives distributions with similar peaks for the three regions, but a larger fraction of ‘fast axons’ for the mid body segment. We simulate the LFP signal that would arise from the distribution of latencies for a 10cm long axon (Fig. 5ce). We find the genu display the most delayed peak. Next, we plot the LFP and latency distribution given the tract lengths calculated from tractography (Fig. 5df). The average tract length for occipital (11.5 cm), motor (9.2 cm) and anterior-frontal (8 cm) fiber tracts were used for the LFP simulation of the splenium, mid-body, and genu, respectively. Using the tractography fibers length estimates we find the splenium is estimated to have the most delayed response, while the genu and motor body have similar conduction delays. This result highlights the potential differences between conduction velocity and delay time.

## 5. Discussion

In this study we provide a biophysical framework which relates qMRI measurements to white matter signal conduction, in an attempt to provide a link between structure features of axons and electrophysiology. We used this framework to test whether the MRI measurement of g-ratio can be used to model changes in conduction properties with age. We estimated the conduction velocity and conduction time in the corpus callosum of healthy human subjects. We found little to no significant difference between the young and old subject in the *g*-weighted measurement, and in conduction estimates in the occipital frontal and motor callosal fibers.

It would have been of great interest to evaluate the sex differences in changes in conduction with age. Unfortunately, we do not have enough data to support such an analysis. Nevertheless, in a previous study we measured g-ratio in the corpus callosum in 80 people of ages 8 to 81 years old. We found no differences in the males and females. Similar results were found by Cercignani et al. (74), using a different dataset, and slightly different measurements to estimate g-ratio.

The biophysical interpretation of the qMRI measurements relates them to properties of signal conduction along a myelinated axon. That is, estimating conduction velocity as a function of axon properties (axonal diameter, myelin thickness, and fiber length). There are other models that could have been used here. For example, given that the conduction velocity is indeed mostly affected by the geometric characteristics of the internode (rather than the node), one can reduce the conduction velocity of the axon to the conduction velocity of the internode. Assuming a semi-infinite cylinder, the conduction velocity is a function of the membrane time constant (*τ*) and length constant (*λ*) (108,109) *θ* = 2*λ*/*τ*. This solution describes unmyelinated axons better than myelinated ones, but it is convenient since it doesn’t require a full simulation of the axon propagation. Different models will produce different predictions of conduction latency, but unless one chooses a model that is based on a very different set of assumptions of the axon biophysics, we do not expect it will lead to a great difference in the results.

When applying a biophysical model to MRI data, it is necessary to extract summary statistics of the microstructure parameters. One possibility is to use the voxels in a certain region (e.g. their mean). To sample white matter pathways, it is advantageous to rely on the tractography results, and either use the tract core values, or obtain the characteristic value for each streamline. While the tract core values are probably less susceptible to partial volume effects, using the streamlines allows an interesting and potentially informative sampling of the space.

Using the streamlines as a way to estimate a distribution of axonal values is a very coarse reduction of the anatomy. Whether or not it is valid depends on the homogeneity of the axons in space within a fiber tract. Future testing is needed to decide if ERP characteristics such as latency or width are helpful when interpreting the structural MRI data. To our knowledge, there is not enough evidence today to determine whether it is helpful to use such analysis. Nevertheless, in our data, the LFP analysis predicts group differences both in the peak time and importantly, in the width of the ERP, which is not available when using the tract core values. The difference in ERP width suggests that the LFP simulation captures somewhat different information in the data than using the tract core values.

Using the LFP analysis, we find a significant difference in conduction delays in the motor region (Fig. 4c). However, it is a very small difference (~2ms), and it is not replicated using the core values (Fig. 3c), suggesting that this result is model-dependent. The existing literature on callosal delays does suggest an elongation of signal transfer time with age (33,34). There are several possible explanations for this discrepancy. First, given that there is an increase in inter-hemispheric transfer time (IHTT) with age, it is possible that the increase is a result of cortical processing and not changes in white matter conduction. Second, as previously discussed (27), the myelin measure used here (MTV) is sensitive, but not specific to myelin. Therefore, as the tissue undergoes changes in composition with age, it is possible that our *g*-weighted measurement is becoming less specific to myelin (i.e., affected by the contribution of non-myelin tissue component). On a similar note, it is possible that the single axon model is too coarse, and it might be necessary to formulate a model that considers the entire axonal populations that affects the MRI measures. Furthermore, while the fibers in the corpus callosum are considered to be coherent in orientation, there is evidence suggesting fine dispersion of the fiber orientation (110). Such variation in the fiber orientation could both introduce error to the estimation of the NODDI parameters, and to the tractography and thus the calculation of tract length.

Third, the g-ratio might not capture enough of the variance that is essential for the calculation of conduction velocity. If the tissue changes with age, it is unlikely that only one aspect of the tissue changes. Unfortunately, the biophysical model is relying on structural parameters that are not available *in vivo*. Therefore, it is possible that one of the parameters we kept constant in the calculation of conduction velocity, in fact changes with age. Simulation studies have shown that the biggest factor of conduction velocity is the geometry of the internode (42,48,111). The internode length, for example, is an important factor that may be modulated and affect conduction velocity (86,87). While we do not have access to it, it was found to be correlated with axon diameter, and we use this relationship to reduce the number of fixed parameters.

It is well established that the axon diameter is a very important determinant of conduction velocity (18,48), and this could have further implication for neural computation (112). Furthermore, studies have shown that the axon diameter distributions are related to information transfer rate and its effectiveness (113,114). Therefore, by fixing the axon diameter we are probably losing information (92,93,95,96). A second consequence of choosing a certain axon diameter (and internode length), is that it changes the effect of g-ratio on the conduction velocity. The lower the axon diameter, the less the g-ratio will affect the velocity prediction. Thus, when calculating the latency (rather than velocity), choosing a small axon diameter will weight the tract lengths more strongly. Our results highlight the significance of the ongoing efforts in the qMRI community to improve and simplify *in vivo* measurement of axon diameter (e.g. 24,49,50,52,102).

Due to the importance of the axon diameter we chose empirically based values, from post mortem histology of human corpus callosum. We chose to fix the axon diameter in the occipital, motor and anterior-frontal tracts of the corpus callosum, to be 1.72, 2.81 and 1.3 *μm*, respectively. The difference in axon diameter values leads to large differences in the conduction estimates between tracts. We found that using the empirically based values of axon diameter indeed produces a range of conduction latencies which are close to those found in the literature. Nevertheless, regarding the g-ratio measurement of the different tracts, we find that in this study the between-tract comparisons largely depend on our sampling strategy. In Supp. Fig. 2 it’s clear that the g-ratio changes drastically along the occipital tract. It is possible that the variability in g-ratio along the occipital tract is due to the other fiber tracts it crosses paths with. The optic radiation, which is adjacent to the splenium fibers (117) and could therefore lead to partial volume effects, is known to be highly myelinated (118) causing a large decrease in g-ratio. The idea that the trend of g-ratio along the occipital callosal fiber is influenced by adjacent fibers is in line with the fact that it is similar between younger and older subjects (Supp. Fig. 2).

Our results point to interesting differences between the three callosal tracts used in this study. We find the g-ratio is the highest in the occipital tract, replicating our previous study (119), as well as other studies (20,29). To estimate the conduction velocity we fix the axon diameter using the weighted average of the axon diameter distributions measured by Aboitiz et al. (89). The mid-body, corresponding to the motor tract, has a large fraction of large axon. This in turn, leads to the faster estimates of conduction velocity and shorter delay time (Fig. 4bd, and Fig. 5c.). Using the entire axon diameter distribution to estimate the conduction properties of the tracts (Fig. 6), we find that given a constant tract length, the splenium and mid-body have very similar latencies, and the genu has the largest delay (Fig. 6e). However, taking into account the tract lengths as measured with dMRI, we find the genu and mid-body display similar latency, while the splenium is the slowest (Fig. 6f). Interestingly, in Fig. 4, we also find that the velocity might change slightly with age, yet the latency does not. Highlighting the fact that while the velocities along the corpus callosum may vary, the latencies could show a different trend as function of the tract length.

The analysis incorporating the axon diameter distributions estimates the slowest conduction will be in the splenium of the corpus callosum, containing occipital fibers (Fig. 6ef). We find the same results using the weighted average of the distributions and the g-ratio measured using MRI (Fig. 4d, and Fig. 5c). Nevertheless, while the former analysis finds similar latencies for the genu and mid-body, the latter estimates the mid-body to have faster conducting fibers. The discrepancy is likely due to the difference between using the entire distribution, and using a summary statistic to represent it: the large fraction of fast axons in the mid-body increases its weighted average dramatically, leading to large differences in the velocity estimates of the three tracts. However, when using the entire axon diameter distribution, we derive distributions of velocities with fairly similar peaks, leading to closer estimated of LFP peak time. Other differences originate from the way the latency distributions are constructed. The latency distributions (from which the LFP is simulated) derived in Fig. 5ac reflects the variance of g-ratio within tract, whereas in Fig. 6 they represent the variance of axon diameters, measured with histology. One last important distinction is that in the axon diameter distribution analysis we choose g-ratio as a function of the axon diameter. This causes larger axons to have larger g-ratio, meaning less myelin. This could decrease the differences between the tracts. Note that while one can assume a certain g-ratio per axon diameter, the opposite deduction is harder to make. This is due to the shape of their relationship: a large range of axon diameter can have similar g-ratios. Furthermore, we do not know how the axon diameter distribution changes with age. These results suggest that more precise histological analysis of axon diameter and g-ratio in the corpus callosum body is needed.

Finally, the literature describing an increase in IHTT with age, did not measure callosal conduction directly, but instead used indirect behavioral measures such as response time. Therefore, it is possible that a more direct estimation of callosal delays will reveal that they are indeed stable with age. The conduction delays can be estimated from the evoked response potential which is measured *in vivo* with methods with high temporal resolution (e.g., ECoG, EEG and MEG). The M/EEG signal represents a conglomeration of many different neural sources of activity (120). Embedded in the signal are the neural responses associated with specific sensory, cognitive, and motor events. Conduction along the corpus callosum can be accessed by presenting subjects with unilateral stimuli (tactile or visual), since such stimuli are first processed in the contralateral hemisphere, and the information is then transferred to the ipsilateral hemisphere via the corpus callosum. By measuring the same early ERP components (e.g., parietal P100) from the different hemispheres, and computing the difference between their time of occurrence, one can estimate the IHTT – the time it took the signal to travel from one hemisphere to the other, which is an estimation of conduction delay along callosal fibers (121,122).

We tested the power of a *g*-weighted MRI measurement to predict conduction delays with age, and we propose that future studies would benefit from testing the framework by measuring conduction delays and structural MRI together. It should be noted, however, that the structural MRI measurements cannot be expected to fully explain the variance in the ERP latency. Of course, a higher resolution and SNR might improve the prediction. Nevertheless, predicting conduction delays can also have other limiting factors. Currently it is impossible to measure key properties, such as the internode length, that affect the conduction velocity. Furthermore, the ERP latency is affected by more than the conduction in white matter. The ERP could be affected by processing time (cortical, or sensory), and potentially some properties of the skull and gyrification that could filter the signal. While the inter-individual variance of such properties is still unclear, it is unlikely to be zero. Therefore, one cannot expect to fully explain the data.

Despite the aforementioned challenges, it is important to conceptualize the relationship between structural MRI measurements and functional EEG recordings using a biophysical framework. We hope that this work can be used as a first step in bringing these two fields together in an effort to explain common sources for individual differences.

## 6. Summary

We provide a framework that relates MRI measurements of averaged microstructural features, to a single axon conduction model, in order to be able to predict white matter conduction *in vivo.* This is a critical step in the process of relating structural MRI measurements to white matter function, in a biologically relevant manner. Such models are highly reductive when we use *in vivo* measurements that are averaged over a large number of axons. Our results further suggest that using *g*-weighted measurement might not be enough to evaluate meaningful differences in conduction with age. It will be important to take the best available qMRI data (axon diameter estimates included), and a complementary measurement of conduction to show to what extent the models are useful. We hope that as the qMRI models develop further, this sort of study will reveal more about the human brain and behavior in healthy development, aging, and dysfunction.

## Acknowledgments

This work was supported by the Ministry of Science Technology & Space, Israel (grant no. 3-13395) awarded to S.B. It was also supported by the ISF Grant (no. 0399306) and the NSF/SBE-BSF Grants (NSF no. 1551330 and BSF no. 2015608) awarded to A.A.M., and a seed grant from the Eric Roland Fund for Interdisciplinary Research administered by ELSC, awarded to A.A.M. and S.B.

We thank Hiromasa Takemura, Mickey London, Leon Deouel, and Gal Atlan for their helpful comments on this manuscript.

---

**Supplementary Figure 1:**
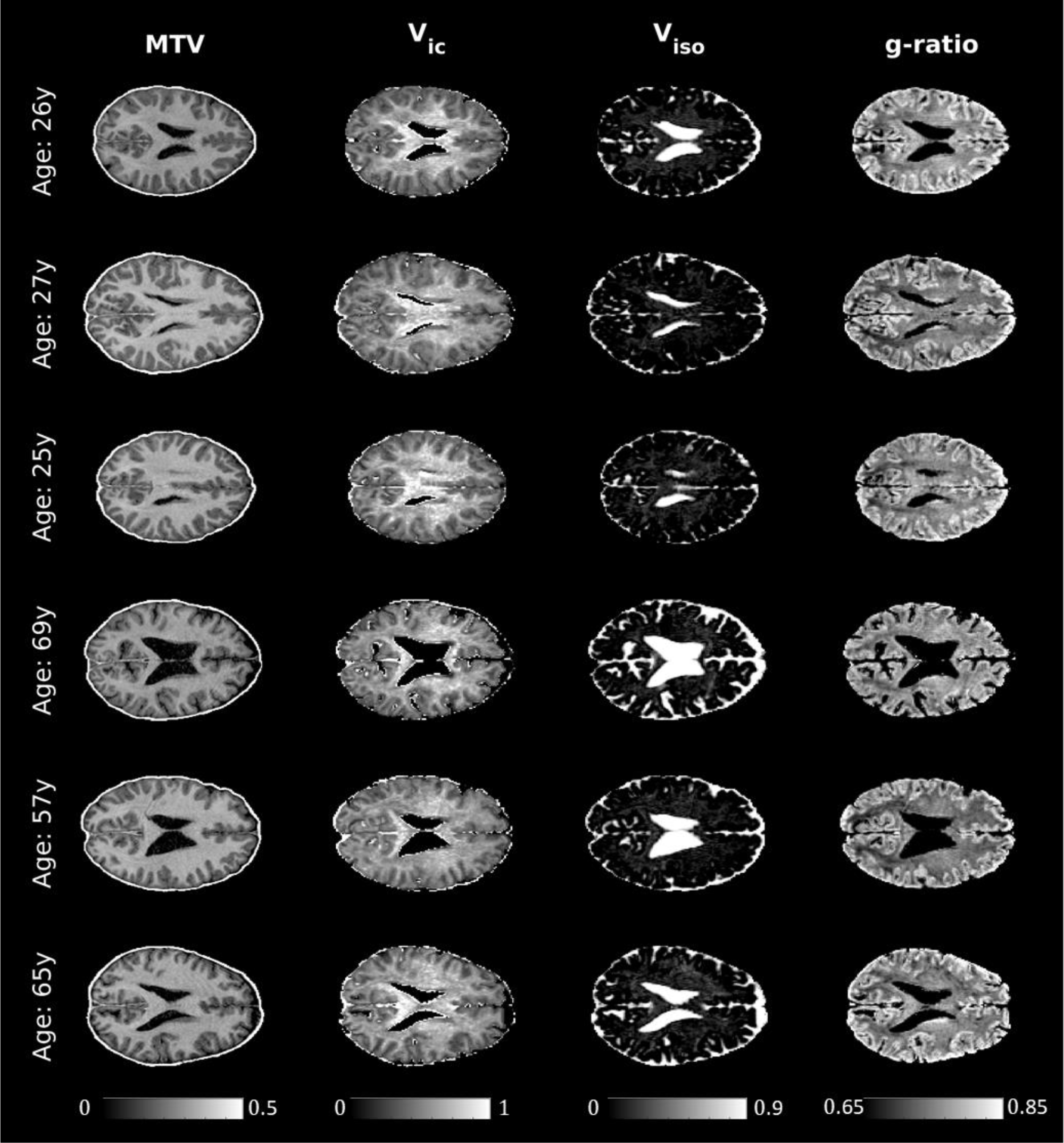
whole brain g-ratio maps. Examples of the qMRI maps used in this study, for 3 younger and 3 older subjects. The maps presented here are the MTV (the non-water fraction), the intracellular and isotropic volume fraction (*V*_*ic*_ and *V*_*iso*_, respectively), calculated as parameters in the NODDI model. The g-ratio is calculated from *MTV*, *V*_*ic*_ *V*_*iso*_.

**Supplementary Figure 2:**
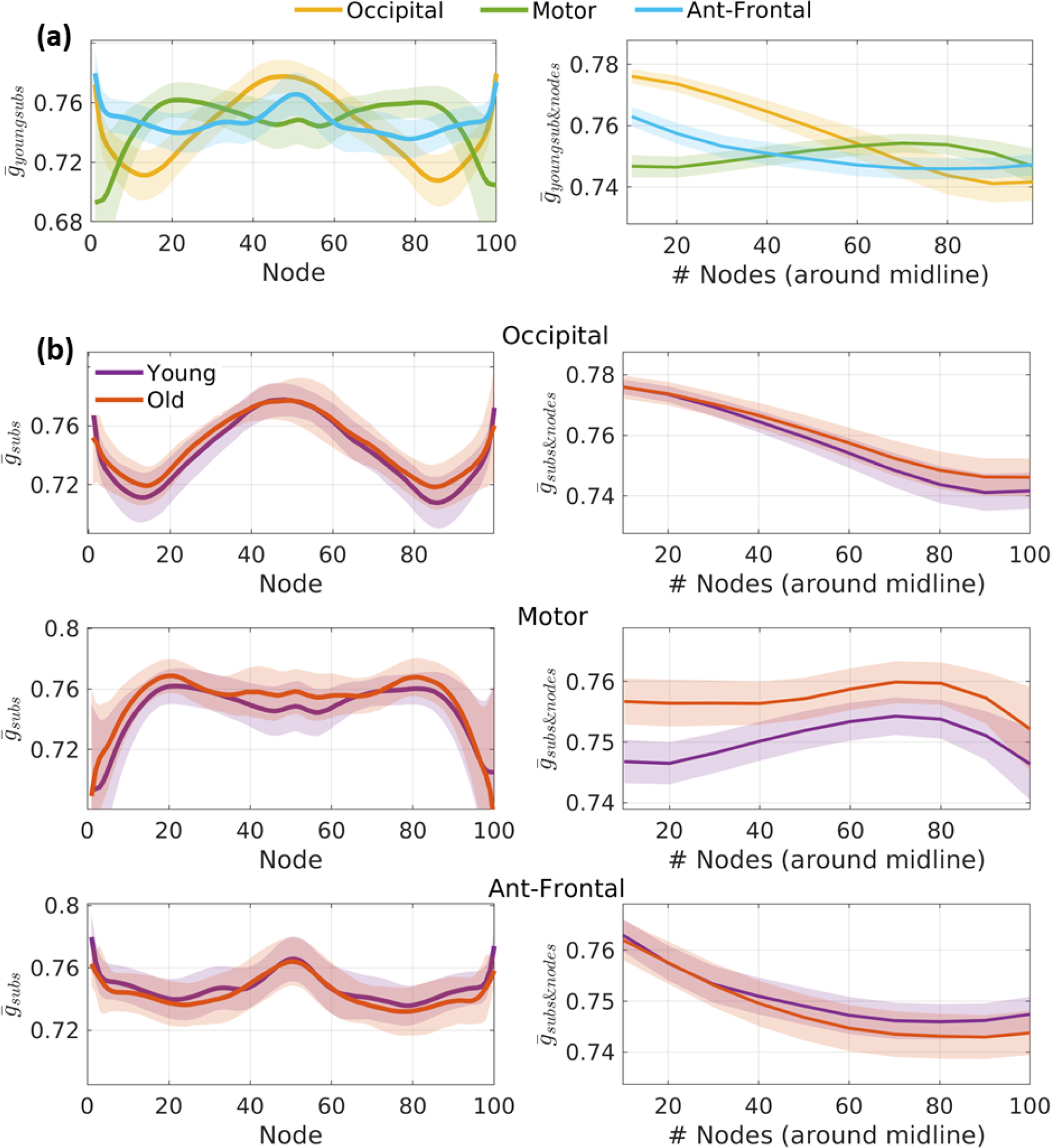
Variance along the fiber tract. **(a)** The left plot shows the average g across younger subjects at each node along the fiber tract (100 nodes, from left to right). The shaded area represents the STD at each node. The three tracts show different relationships with respect to the node location. To characterize the g of a tract, we average over several nodes. The right plot is the average as a function of the number of nodes taken around the midline (node 50), from 10 nodes to the entire tract (shaded area represents STD). The relationship between the callosal tracts changes as a function of the number of nodes taken to calculate the mean. **(b)** The plots in (a) are shown separately for the different fiber tracts, comparing the average g of younger (purple) and older (orange) subjects. While there are slight variations in the age-group differences as a function of node-number, the overall relationship remains the same.

**Supplementary Figure 3:**
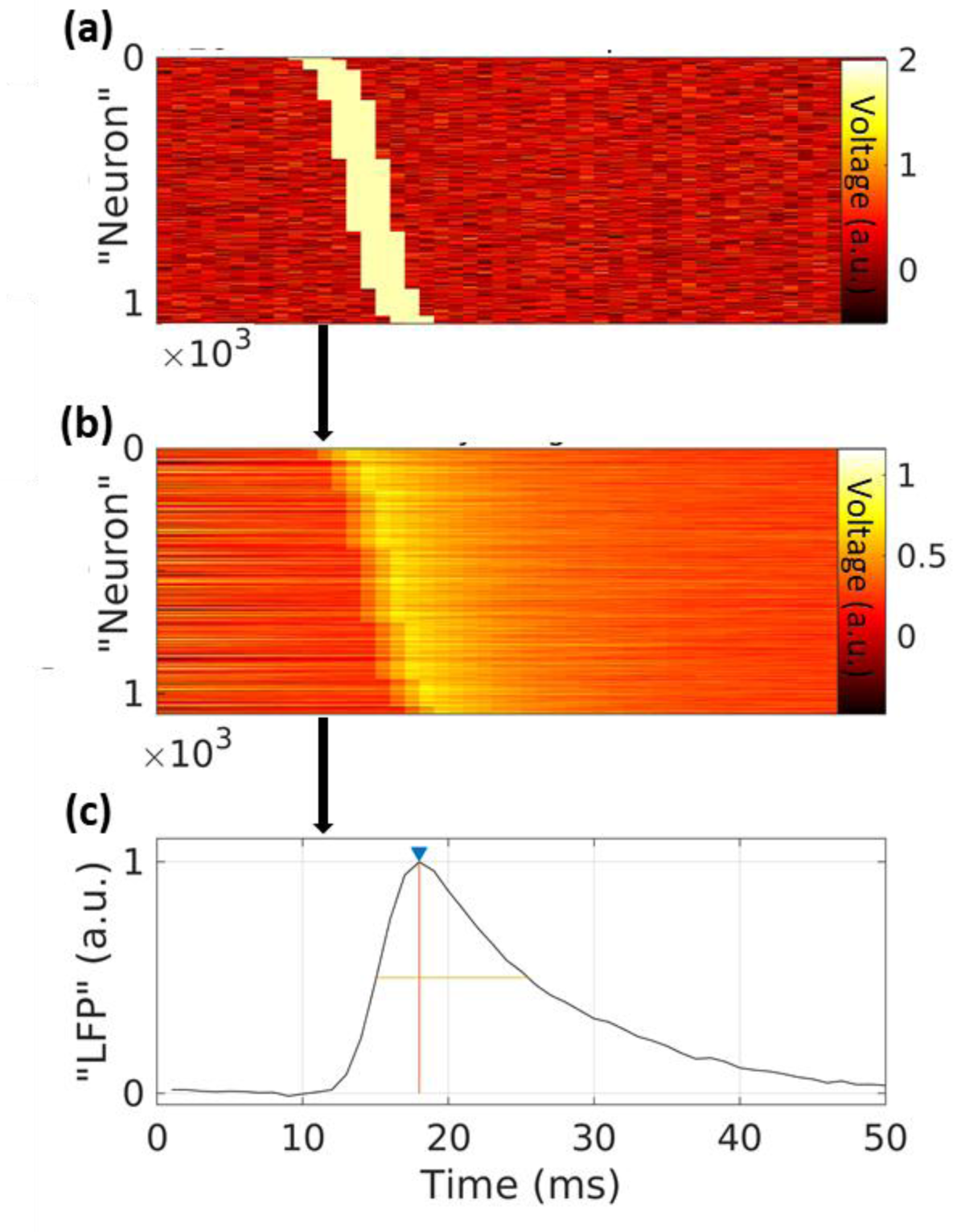
LFP simulation. The distribution of latencies, calculated from the streamlines’ g-ratio and length, is used as input for the simulation of leaky-integrating neurons. **(a)** The input is modeled as Gaussian noise, with a delta function of the corresponding latency of the streamline. **(b)** The input is then leaky-integrated over time. **(c)** The LFP is the sum over the integrated time series. The signal is normalized to have a peak of 1 [a.u]. Finally, we calculated the peak time and the full width half maximum of the ERP.

